# Engineering ecologically complementary rhizosphere probiotics using consortia of specialized bacterial mutants

**DOI:** 10.1101/2022.03.16.484597

**Authors:** Chunlan Yang, Jingxuan Li, Alexandre Jousset, Xiaofang Wang, Zhihui Xu, Tianjie Yang, Xinlan Mei, Zengtao Zhong, Yangchun Xu, Qirong Shen, Zhong Wei, Ville-Petri Friman

## Abstract

While bacterial diversity is beneficial for the functioning of rhizosphere microbiomes, multi-species bioinoculants often fail to promote plant growth. One potential reason for this is that competition between inoculated consortia members create conflicts for their survival and functioning. To circumvent this, we used transposon mutagenesis to increase the functional diversity within *Bacillus amyloliquefaciens* bacterial species and tested if we could improve plant growth-promotion by assembling consortia of closely related but functionally specialized mutants. While most insertion mutations were harmful, some improved strains’ plant growth-promotion traits without increasing antagonism between them. Crucially, plant growth-promotion could be improved by applying these specialist mutants as consortia, leading to clear positive relationships between consortia richness, plant root colonization and protection from bacterial wilt disease. Together, our results suggest that increasing intra-species diversity could be an effective way to increase probiotic consortia multifunctionality, leading to more stable plant growth-promotion throughout growth cycle via insurance effects.

## Introduction

Bacterial rhizosphere microbiome diversity has been strongly associated with beneficial effects on plant growth in both natural and agricultural environments (1, 2). Such beneficial diversity effects are often associated with increased functional diversity, where different taxa have specialist roles in plant growth-promotion that can include nutrient solubilization, plant immune priming or interactions with other beneficial or pathogenic rhizosphere microbes (3, 4). While several studies have tried to harness positive effects of bacterial diversity and multifunctionality for the crop production (5–7), diverse inoculants often fail to produce the desired benefits in field conditions. This discrepancy could rise due to constraints set by external factors such as resource availability (8–10), that constrains the expression of plant growth-promotion traits, or the presence of competitors, parasites and predators that could reduce the survival of the inoculated bacteria in the rhizosphere (11, 12). It is also possible that interactions between probiotic consortia members could affect their survival in the rhizosphere if they show antagonism towards each other (13–15). As antagonistic traits often increase the invasion success of probiotic bacteria, it has thus been suggested that inoculant design should aim to minimize negative interactions within consortia, while sustaining antagonism between pathogens or resident microbiota (15–17). One way to achieve this is to use community ecological theory to design consortia based on niche complementarity in order to limit competition while still promoting potential facilitative interactions between inoculant members (16, 17).

Ecological complementarity and division of labor can efficiently reduce competition between coexisting species or genotypes by allowing different community members to use different niches or specialize to different functions within the same niches (18, 19). In multi-species communities, ecological complementarity could be achieved via community assembly, where competitively similar species exclude each other, leading to coexistence of niche specialists (20). In monocultures, ecological complementarity could evolve via character displacement, where different genotypes adapt to use specific environmental niches (19, 21, 22). Ecological complementarity could lead to improved ecosystem functioning if productivity is higher in consortia than in associated monocultures of individual species or genotypes (23, 24). Such overperformance (or overyielding) could be driven by different mechanisms, including resource partitioning, abiotic facilitation, or biotic feedbacks (25). In contrast, division of labor could improve consortia functioning even within a single niche via specialization into different tasks, as has been demonstrated with *Bacillus subtilis* bacterium during biofilm matrix production (18). Division of labor however also requires that interactions between community members are cooperative, and that all individuals gain an inclusive fitness benefit from the interaction (26), which could make communities less stable. In support for this, it has been shown that genetic division of labor between specialized *B. subtilis* mutants can collapse at the evolutionary timescale, where further adaptation leads to ecological autonomy of interacting consortia members (27). In addition to occupying different niches, or performing specialized tasks, the consortia coexistence could be favored due to facilitative interactions (16, 28) via cross-feeding, where bacterial community members might benefit from each other’s secretions (29), or cooperation where genetically related individuals together perform different functions, either altruistically or mutualistically (30). Theoretical work has suggested that facilitative interactions could take place at intermediate levels of genetic mixing, where strains produce only a subset of secretions and rely on other genotypes for the complementary traits (31). However, it has also been predicted that such facilitative interactions might be inefficient, leading to reduced productivity by specialist consortia compared to the generalist genotypes (31). It thus remains unclear how to best optimize consortia multifunctionality in probiotic inoculant design, and whether this should be based on ecological complementarity, division of labor or facilitation between consortia members.

Here, we focused on studying if increasing intra-species diversity of a single bacterium could offer a viable strategy for improving consortia multifunctionality without introducing conflicts between the consortia members. Consortia diversity has been linked to multifunctionality (32–36), beneficial insurance effects (37), emergent consortia-level properties via increased production of antimicrobials (38, 39) or intensified resource competition(40, 41), and positive species identity effects linked with plant health (15, 42–45). Most studies have reported these diversity effects in multi-species communities, neglecting the potential benefits of intra-species diversity, which has also been shown to play an important role in the functioning of microbial ecosystems (18, 24, 46–49). To study this, we used a combination of synthetic biology and biodiversity-ecosystem functioning theory (32, 37) to harness the benefits of intra-species diversity of *Bacillus amyloliquefaciens* T-5 bacterium for plant growth-promotion. We chose to focus on *B. amyloliquefaciens* T-5 strain because, it originates from the tomato rhizosphere (50) and has been previously shown to protect plants from various diseases, including bacterial wilt, which is caused by phytopathogenic *Ralstonia solanacearum* bacterium (51). To increase functional diversity within a single species, we first created a *B. amyloliquefaciens* mutant library using TnYLB-1 transposon mutagenesis (52) and phenotyped 479 resulting mutants regarding four important plant growth-promotion traits in the lab: biomass production, biofilm formation, swarming motility and pathogen suppression (4, 53, 54). A subset of functionally specialized mutants was then used to design probiotic inoculant consortia with varying richness levels (*i.e*., manipulating the number of mutants per consortium) to directly test if plant growth-promotion could be improved relative to the wild-type strain by using highly related multi-genotype communities. Richness manipulation was chosen to capture potential benefits effects rising due to ecological complementarity, division of labor or facilitation. Using this approach, we aimed to attain following advances. First, we expected that consortia diversity should increase multifunctionality through inclusion of functionally specialized mutants. Second, we expected that mutant consortia consisting of specialists with low niche overlap would show higher functioning due to ecological complementarity, which could even lead to overyielding, *i.e*., better than expected consortia functioning based on the individual performance of its members. Finally, by using a consortium of mutants we hoped to reduce the effect of physiological trade-offs experienced at the individual clone level, potentially via task allocation and division of labor (18, 55–58), and to potentially maintain cooperation within bacterial species due to close genetic relatedness and kin selection (59–62). Together, these mechanisms could provide stability to plant growth-promotion via insurance effects, where diverse probiotic consortia would be more likely to retain their functioning even if some inoculant members fail to survive or perform poorly in the rhizosphere (63). Our results suggest that synthetic biology can be used to increase the functional diversity of a probiotic bacterium without imposing intra-species conflicts, leading to improved root colonization and plant protection by specialized probiotic mutant consortia.

## Materials and methods

### Bacterial strains and culture conditions

We used phytopathogenic *Ralstonia solanacearum* QL-Rs1115 (64) and *Bacillus amyloliquefaciens* T-5 biocontrol (50) strains as our model bacterial species. The *B. amyloliquefaciens* T-5 can suppress the growth of *R. solanacearum* QL-Rs1115 by competing for space and nutrients in the rhizosphere (65) and by producing various antibacterial secondary metabolites (66). Both bacterial stocks were cryopreserved at −80 °C in 30% glycerol stocks. Prior starting the experiments, active cultures were prepared as follows: *B. amyloliquefaciens* T-5 was grown at 37 °C in Lysogeny Broth (LB-Lennox, 10.0 g L^-1^ Tryptone, 5.0 g L^-1^ yeast extract, 5.0 g L^-1^ NaCl, pH=7.0) and *R. solanacearum* QL-Rs1115 was grown at 30 °C in Nutrient Broth (NB, 10.0 g L^-1^ glucose, 5.0 g L^-1^ peptone, 0.5 g L^-1^ yeast extract, 3.0 g L^-1^ beef extract, pH = 7.0) for 24 h.

### Generation of *Bacillus amyloliquefaciens* T-5 transposon mutant library

To increase the intra-species diversity of *B. amyloliquefaciens*, we generated a random transposon insertion mutant library by using a TnYLB-1 transposon derivative, carried in the thermosensitive shuttle plasmid pMarA (*SI Appendix*, Table S1), which was electro-transformed to bacteria as previously described by Zakataeva *et al*. (67, 68). The cells with intact pMarA plasmid contained resistance cassettes to both erythromycin and kanamycin, while the cells with integrated transposons were resistant only to kanamycin. Transposon mutant library was created as follows. An overnight *B. amyloliquefaciens* T-5 cell culture grown in neutral complex medium (NCM, 17.4 g L^-1^ K_2_HPO_4_, 11.6 g L^-1^ NaCl, 5 g L^-1^ glucose, 5 g L^-1^ tryptone, 1 g L^-1^ yeast extract, 0.3 g L^-1^trisodium citrate, 0.05 g L^-1^ MgSO_4_.7H_2_O, and 91.1 g L^-1^ sorbitol, pH = 7.2) was diluted 25-fold with fresh NCM medium supplemented with 5 mg mL^-1^ of glycine and grown at 30 °C for 3 h on a rotary shaker (170 rpm). After 1h incubation (at OD_600_ ~ 0.8), cells were cooled on ice, harvested by centrifugation (8,000 × g for 6 min at 4 °C) and washed four times with ice-cold electrotransformation buffer (ETM, 0.5 M sorbitol, 0.5 M mannitol, and 10% glycerol). Resulting pellets were resuspended in ETM buffer supplemented with 10% PEG 6000 and 1mM MgCl_2_, yielding approximately 10^10^ cells mL^-1^. Cells were then mixed with 500 ng of plasmid DNA in an ice-cold electrotransformation cuvette (2-mm electrode gap), and after 1-3 min incubation at room temperature, exposed to a single electrical pulse using a MicroPulser Electroporator (Bio-Rad Laboratories) at field strength of 7.5 kV cm^-1^ for 4.5-6 ms. Immediately after the electrical discharge, cells were transferred into 1 mL of LB, incubated with gentle shaking at 30 °C for 3-8 h, and plated on LB agar containing 10 μg mL^-1^ erythromycin. Transformants were selected after 36-48 h incubation at 30 °C. To generate final transposon library, erythromycin-resistant colonies with plasmids were individually transferred to fresh LB and incubated overnight at 30 °C, after cultures were diluted, spread on LB plates supplemented with 10 μg mL^-1^ kanamycin, and incubated for 24 h at 46 °C. As the plasmid cannot replicate at 46°C, only cells with an integrated transposons grew and could be separated. A total of 2000 transformed colonies were isolated and individually cryopreserved in 30% glycerol at −80 °C.

### Phenotypic characterization of *B. amyloliquefaciens* T-5 mutant library *in vitro*

The wild-type strain and 479 randomly selected mutants (out of a total of 2000 created mutants) were phenotyped for following plant-growth promoting traits: swarming motility, biomass production, biofilm formation and pathogen suppression via production of antibiotics. These traits were selected due to their known importance for *B. amyloliquefaciens* competitiveness in the rhizosphere and their involvement in pathogen suppression (69–72). To prepare bacterial inoculants, frozen colonies were picked and pre-grown overnight in LB at 37 °C, washed three times in 0.85% NaCl and adjusted to density of 10^8^ cells mL^-1^. In addition to each individual trait, we also calculated the average of all measured traits and used the resulting ‘monoculture average performance’ (73) index to compare mutants’ overall performance (*SI Appendix*, Dataset S1).

#### Swarming motility

was measured using a previous method described by Kearns (74). Briefly, a drop (2 μL equaling 10^6^ cells mL^-1^) of each *B. amyloliquefaciens* mutant was inoculated into the center of 0.7% agar LB plates supplemented with 10 μg mL^-1^ of kanamycin. After 24 h incubation at 30 °C, swarming motility was evaluated as the radius of the colony. Three replicates were used for each mutant.

#### Biomass production

was assessed on 96-wells microtiter plates (at 30 °C with agitation) in 200 μL of ‘recomposed exudate’ medium (abbreviated as ‘RE’, which contained: 0.5 g L^-1^ MgSO_4_.7H_2_O, 1.0 g L^-1^ K_2_HPO_4_, 0.5 g L^-1^ KCl, 1.0 g L^-1^ yeast extract, 1.2 mg L^-1^ Fe_2_(SO_4_)_3_, 0.4 mg L^-1^ MnSO_4_, 1.6 mg L^-1^ CuSO_4_, 2 g L^-1^ (NH_4_)_2_SO_4_, 0.8 g L^-1^ glucose, 1.3 g L^-1^ fructose, 0.2 g L^-1^ maltose, 0.02 g L^-1^ ribose, 5.6 g L^-1^ citrate, 1.4 g L^-1^ succinate, 0.2 g L^-1^malate, 0.8 g L^-1^ casamino acids (75)). The initial inoculum size for each mutant was adjusted to an optical density (OD_600_) of 0.05 (equalling ^~^10^5^ cells mL^-1^). After 24 h of growth, OD_600_ was measured as a proxy for biomass production. Three replicates were used for each mutant.

#### Biofilm formation

was assessed as described previously (76) using 24-well microtiter plates instead of 96-well plates. Briefly, 10 μL (10^6^ cells mL^-1^) of *B. amyloliquefaciens* cells were added into 1 mL of biofilm-promoting growth medium (MSgg minimal medium: 2.5 mM PBS [pH 7.0], 100 mM MOPS [pH 7.0], 50 μM FeCl_3_, 2 mM MgCl_2_, 50 μM MnCl_2_, 1 μM ZnCl_2_, 2 μM thiamine, 50 mg phenylalanine, 0.5% glycerol, 0.5% glutamate and 0.7 mM CaCl_2_) on 24-well polyvinyl chloride microtiter plates and incubated without agitation for 24 h at 30 °C (77). The growth medium and planktonic cells were removed, and remaining cells adhered on well walls were stained with 1% crystal violet dissolved in washing buffer (0.15 M (NH_4_)_2_SO_4_, 100 mM K_2_HPO_4_ [pH 7], 34 mM sodium citrate, and 1 mM MgSO_4_) for 20 mins at room temperature. Plates were then rinsed with demineralized water to remove excess crystal violet, after the remaining crystal violet bound to well wall biofilms were solubilized in 200 μL of solvent (80% ethanol, 20% acetone). Biofilm formation was defined as the optical density of crystal violet at OD_570_. Three replicates were used for each mutant.

#### Pathogen suppression

via production of antibiotics was assessed as inhibition of *R. solanacearum* QL-Rs1115 strain using an agar overlay assay (78). Briefly, small volume drops (2 μL) of each *B. amyloliquefaciens* mutant and wild-type strain were spotted on NA plates and incubated for 24 h at 30 °C. Plates were then chloroform-fumigated to kill all the bacteria (78), leaving only the secreted antimicrobials on the agar medium and fully covered with soft agar overlay containing *R. solanacearum* suspension (with a final concentration of approximately 10^7^ cells mL^-1^). The pathogen suppression of each mutant was defined as the area of the *R. solanacearum* inhibition halo around the *B. amyloliquefaciens* colony (in mm^2^), which is proportional to antibiotic production (79). Three replicates were used for each mutant.

### Selecting a representative subset of *B. amyloliquefaciens* T-5 mutants for greenhouse experiments

In order to select a representative subset of mutants for greenhouse experiments, K-means clustering (80) was used to visualize trait differences between the wild-type and 479 characterized mutants based on swarming motility, biomass production, biofilm formation and pathogen suppression (*SI Appendix*, Dataset S1). Mutants clustered to three clear groups based on the first two principal components, which explained 57% and 20.5% of the total variation, respectively. We randomly selected approximately 10% of strains from each cluster, resulting in a subset of 47 mutants, which were used for greenhouse experiments (26, 11 and 10 mutants from Clusters 1, 2 and 3, respectively, *SI Appendix*, Dataset S2). These 47 mutants were further analyzed to determine the disrupted genes by TnYLB-1 transposon insertion using the Inverse Polymerase Chain Reaction (IPCR) method as previously described by Le Breton and colleagues (52). First, 5 μg of genomic DNA isolated from each respective transposon mutant was digested with Taq I and circularized using ‘Rapid Ligation’ kit (Fermentas, Germany). IPCR was carried out with ligated DNA (100 ng), using oIPCR1 and oIPCR2 primers (*SI Appendix*, Table S2). The cloned sequences were then purified using PCR purification kit (Axygen, UK) and the flanking genomic regions surrounding the transposon insertion sites were sequenced using the primer oIPCR3 (*SI Appendix*, Table S2). Obtained DNA sequences were compared against available databases (GenBank and *Bacillus* Genome Data-base) using the BLASTX and BLASTN (81) available at the NCBI, and against the complete ancestral *B. amyloliquefaciens* T-5 genome sequence (Accession: CP061168, *SI Appendix*, Fig. S1A, Dataset S2). The functional classification of disrupted genes for all 47 transposon mutants is summarized in *SI Appendix*, Fig. S1B.

### Assessing the performance of individual *B. amyloliquefaciens* T-5 mutants in a greenhouse experiment

All selected 47 mutants and the wild-type strain were individually screened for their ability to colonize tomato rhizosphere and protect plants against infection by *R. solanacearum* QL-Rs1115 pathogen strain in a 50-day long greenhouse experiment. Surface-sterilized tomato seeds (*Lycopersicum esculentum, cultivar “Jiangshu”*) were germinated on water agar plates in the dark at 28 °C for 2 days, before sowing to sterile pots containing wet vermiculite (Huainong, Huaian soil and fertilizer Institute, Huaian, China). Ten-days old tomato seedlings (at three-leaves stage) were then transplanted to seedling trays containing natural, non-sterile soil collected from a tomato field in Qilin Town, Nanjing, China (64). Plants were inoculated with individual *B. amyloliquefaciens* mutants by drenching, resulting in a final concentration of 10^7^ colony forming units (CFU) g^-1^ soil (82). The *R. solanacearum* strain was inoculated using the same method one week later at a final concentration of 10^6^ CFU g^-1^ soil. Positive control plants were treated only with *R. solanacearum*, while negative control plants received no bacterial inoculants. Three replicated trays were set up for each treatment, with 20 seedlings (in individual cells) per tray. Each tray was considered as one biological replicate. Tomato plants were grown for 30 days after pathogen inoculation with natural temperature (ranging from 25°C to 35°C) and lighting variation (around 16 h of light and 8 h of dark). Seedling trays were rearranged randomly every second day and regularly watered with sterile water.

### Quantifying *B. amyloliquefaciens* mutants’ root colonization and plant protection in the rhizosphere

The root colonization and plant protection of 47 *B. amyloliquefaciens* T-5 mutants was quantified individually as a change in their population densities in the tomato rhizosphere after 5, 15 and 30 days of *R. solanacearum* pathogen inoculation (days post pathogen inoculation, i.e., ‘dpi’). At each sampling time point, three independent plants per inoculated mutant were randomly selected and sampled destructively by carefully uprooting the plant and gently removing the soil from the root system by shaking. After determining plant fresh weight, the root system of each plant was thoroughly ground in 5 mL of 10 mM MgSO_4_·7H_2_O using a mortar, and serial dilutions of root macerates were plated on a semi-selective *Bacillus* medium (83) consisting of 326 ml L^-1^ vegetable juice (V8, Campbell Soup Co., USA), 33 g L^-1^ NaCl, 0.8 g L^-1^ dextrose, 16 g L^-1^ agar (pH 5.2, adjusted with NaOH) supplemented with 45 mg L^-1^ cycloheximide and 22.5 mg L^-1^ polymyxin B (84). This media was used to count the densities of *B. amyloliquefaciens* T-5 wild-type, and the same media supplemented with 10 μg mL^-1^ kanamycin was used to count the densities of mutant strains. Plates were incubated at 30 °C for 30 h and bacterial densities expressed as CFU per gram of root biomass. The effect of *B. amyloliquefaciens* wild-type and mutants on plant protection was measured as the reduction of bacterial wilt disease symptoms during the experiment (based on the proportion of plants showing wilting symptoms). The first wilting symptoms appeared 7 dpi and the proportion of diseased plants quantified at 5, 15 and 30 dpi were used in analyses. Plant protection was expressed as the relative reduction in the number of wilted plants compared to the positive control (only *R. solanacearum* inoculated in the absence of *B. amyloliquefaciens* T-5 mutants or wild-type).

### Assembly of functionally complementary *B. amyloliquefaciens* mutant consortia

To test if the performance of *B. amyloliquefaciens* T-5 mutants could be improved by using consortia of functionally complementary mutants, a subset of eight best-performing mutants excelling at different phenotypic traits were selected (*SI Appendix*, Table S3, Table S4, Dataset S2). Specifically, these included two mutants that showed high swarming motility (M108: *parE*, M124: *DeoR*), high biomass production (M59: *comQ*, M109: *hutI*), biofilm formation (M54: *hutU*, M143: *YsnB*), and slightly improved pathogen suppression (M38: *nhaC*, M78: *dfnG; SI Appendix*, Table S3, Table S4) relative to the wild-type strain. None of the mutants showed antagonistic effects towards each other based on agar overlay assays. These eight mutants were then used to assemble a total of 29 consortia with 2, 4 or 8 mutants following a substitutive design where each mutant was equally often present at each richness level (see Table S5 for detailed consortia assembly). Mutants were mixed in equal proportions in each consortium with final total bacterial density of 10^8^ cells mL^-1^ (e.g., 50:50% or 25:25:25:25% in two and four mutant consortia, respectively). This design has previously been used to investigate biodiversity-ecosystem functioning relationships in plant-associated bacterial communities (37, 85), allowing disentangling the effects due to consortia richness, composition and mutant strain identity. All mutants were also tested individually and the same 29 mutant consortia were used for both *in vitro* lab and *in vivo* greenhouse experiments.

### Phenotypic characterization of *B. amyloliquefaciens* consortia performance *in vitro* and consortia root colonization and plant protection in the tomato rhizosphere

The performance of each mutant and assembled consortium was assessed *in vitro* in the lab by measuring traits as mono- and co-cultures following the same methods as described previously (swarming motility, biomass production, biofilm formation and pathogen suppression). Mutant strains were prepared individually from frozen stocks by growing overnight in liquid LB, pelleted by centrifugation (4,000 × g, 3 min), washed three times with 0.85% NaCl and adjusted to a density of 10^8^ cells mL^-1^. Consortia were then assembled following the substitutive design describe earlier (*SI Appendix*, Table S5) by mixing mutants in equal proportions for each consortium with final total bacterial density of 10^8^ cells mL^-1^ (e.g., 50:50% or 25:25:25:25% in two and four mutant communities, respectively). Consortia traits were characterized as described previously and compared with the ancestral *B. amyloliquefaciens* wild-type strain. The root colonization and plant protection of *B. amyloliquefaciens* T-5 consortia were quantified in greenhouse experiments following previously described methods.

### Statistical analyses

Data were analyzed with a combination of ANOVA, principal component analysis (PCA), linear regression models and unpaired two-sample Wilcoxon tests. Individually measured mutant traits data was normalized between 0 and 1 across the all collection using min-max normalization (86). In addition, the different phenotypic traits were combined into a ‘Monoculture average performance’ index, which was calculated as the mean of the four standardized traits for each mutant. Monoculture average performance and consortia traits values were also min-max normalized between 0 and 1 for subsequent analyses. To classify mutants into different functional groups, K-means algorithm (‘kmeans’ function) was used and clusters were visualized using PCA (‘princomp’ in ‘vegan’ package) based on multivariate trait data. The *B. amyloliquefaciens* T-5 abundance data measured in root colonization assays were log_10_ transformed and disease incidence data were arcsine square root-transformed prior the analyses. Linear regression models were used to explain root colonization and plant protection with mutant traits, monoculture average performance and consortia richness. Mean treatment differences were analyzed using two-sample Wilcoxon test (‘wilcox.test’ function) and analysis of variance (ANOVA, ‘aov’ function). All statistical analyses were performed using R 3.5.2 (R core Development Team, Vienna, Austria).

## Results

### Effects of transposon insertions on *B. amyloliquefaciens* T-5 traits measured *in vitro* and *in vivo*

We first quantified the effects of transposon insertions on *B. amyloliquefaciens* T-5 traits using 479 mutants that were randomly selected across the whole mutant library (2000 mutants in total). Most insertions had negative effects on the four measured traits, with more than half of the mutants showing reduced swarming (58.7%), biomass production (67.2%) and biofilm formation (60.8%) compared to the wild-type strain (Fig. 1A, *SI Appendix*, Dataset S1). In contrast, the median effect of insertions on the pathogen suppression was neutral, and 51.1% of the mutants showed only a moderate increase in their suppressiveness (Fig. 1A, *SI Appendix*, Dataset S1). In line with this finding, the distribution of effects of insertions on each trait was skewed, where beneficial mutations resulted mainly in a moderate improvement, while harmful mutations often led to severe reduction in measured bacterial traits (Fig. 1A). Moreover, several insertions caused trade-offs, where improvement regarding one trait led to a reduction in the expression of other traits (Fig. 1B). For example, swarming motility correlated negatively with biofilm production, while biomass production led to a trade-off with both biofilm production and pathogen suppression (Fig. 1B). These results thus further suggest that transposon insertions constrained the simultaneous expression of multiple traits, leading to specialized *B. amyloliquefaciens* T-5 mutants, which could be clustered in three phenotypically distinct groups (Adonis test: R^2^ = 0.5283, p < 0.001, Fig. 1C). Relative to the wild-type strain, mutants belonging to the cluster 1 showed increase in biofilm formation and pathogen suppression but reduced biomass and no change in biofilm production (Fig. 1D, *SI Appendix*, Table S6). Mutants in the cluster 2 showed improved swarming motility and reduced pathogen suppression but no significant changes in biomass production or biofilm formation (Fig. 1D, *SI Appendix*, Table S6). Finally, mutants grouped in the cluster 3 had poor performance overall, showing highly reduced swarming motility and pathogen suppression, with no changes in biomass production or biofilm formation (Fig. 1D, *SI Appendix*, Table S6). To test if the mutants grouped in different clusters also differed in their tomato root colonization or ability to protect plants from pathogen infections, 47 mutants representing all three clusters were randomly selected for a greenhouse experiment (the specific effects of insertions on biological processes, cellular components and molecular function for all mutants are shown in *SI Appendix*, Figure S1 and Dataset S2). Compared to the wild-type, 57% of *Bacillus* mutants (27/47) reached lower population densities in the rhizosphere (30 days post-pathogen inoculation (dpi)), and this was especially clear for mutants belonging to clusters 2 and 3. In contrast, mutants belonging to the cluster 1 retained efficient root colonization and some of them showed improved root colonization relative to the wild-type. Similarly, while 93% of mutants (44/47) exhibited reduced plant protection relative to the wild-type strain, this was less clear with mutants belonging to cluster 1, whom a few showed even improved plant protection relative to the wild-type (30 dpi, Fig. 1E, F, *SI Appendix*, Table S7). Together, these results suggest that while most transposon mutants showed reduced performance relative to the wild-type strain, some of them showed improvement in at least in one plant growth-promotion trait, which often led to trade-offs with some other traits.

**Figure 1.**
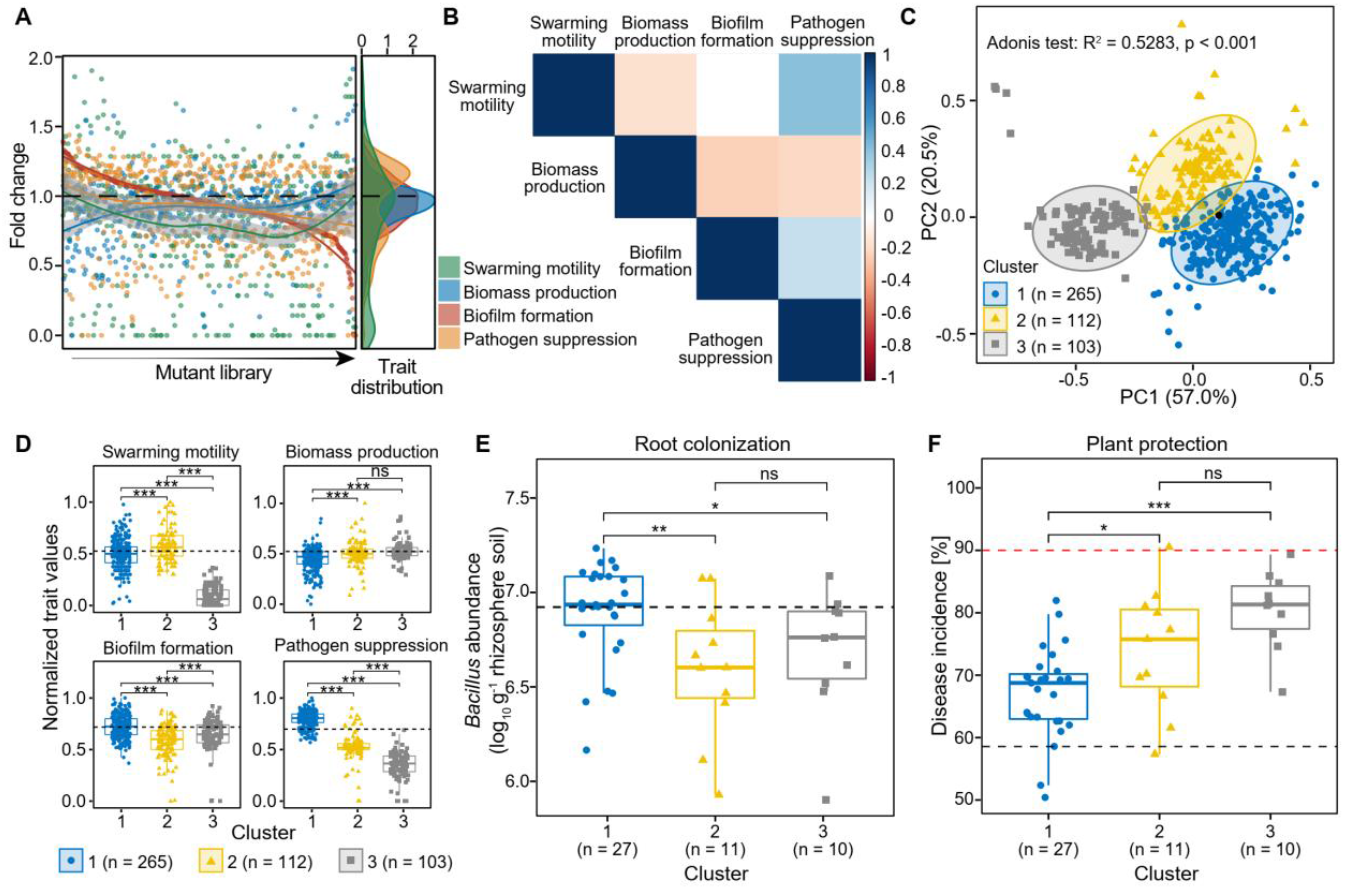
The effects of transposon insertions on the traits of 479 *B. amyloliquefaciens* T-5 mutants measured *in vitro* and *in vivo*. (A) Distribution of fold changes regarding mutants’ swarming motility, biomass production, biofilm formation and pathogen suppression relative to the wild-type strain (dashed line). (B) Pairwise covariance matrix between individual traits, where the red cells indicate negative trait correlations (trade-offs) and the blue cells positive traits correlations; white cells indicate no correlation between the traits. (C) Principal coordinates analysis showing the clustering of all mutants and the wild-type (black point) based on K-means algorithm of four measured traits. (D) Mean trait differences between clusters based on unpaired two-samples Wilcoxon test. (E) Root colonization of 47 *B. amyloliquefaciens* T-5 mutants from cluster 1-3 relative to the wild-type strain (black dashed line) based on cell densities in the root system 30 days post pathogen inoculation (dpi). (F) Plant protection of 47 *B. amyloliquefaciens* T-5 mutants from cluster 1-3 relative to the wild-type strain (black dashed line), and negative ‘pathogen-only’ control (red dashed line), quantified as bacterial wilt disease incidence 30 dpi. Pairwise differences in D-F were analyzed using unpaired two-samples Wilcoxon test: *** denotes for statistical significance at p < 0.001; ** denotes for statistical significance at p < 0.01; * denotes for statistical significance at p < 0.05; ns denote for no significance.

### Designing and testing the performance of mutant consortia *in vitro* and *in vivo*

To test if *B. amyloliquefaciens* T-5 performance could be improved by using mutant consortia, we chose eight mutants that showed higher performance relative to the wild-type strain regarding to one of the plant growth-promotion traits measured *in vitro* (*SI Appendix*, Table S3, Table S4, Dataset S2; two representative mutants per each measured traits selected). We first confirmed that the mutants did not show direct antagonism towards each other using agar overlay assays. As no inhibition was observed, specialist mutants were then used to assemble a total of 37 consortia that varied in their richness level (1, 2, 4, or 8 mutants), following a substitutive design, where each mutant was equally often present at each richness level (see Table S5 for detailed composition of consortia). We hypothesized that consortia could show improved performance due to ecological complementarity where different mutants ‘specialize’ respective to different traits, overcoming trade-offs experienced at individual strain level (Fig. 1B). Alternatively, consortia could show other biodiversity-associated benefits relative to the wild-type such as emergent synergism division of labor or facilitation. We first tested the consortia performance regarding the four traits measured *in vitro*. We found that relative to wild-type strain, only a few consortia showed improved performance regarding swarming motility (8 of 37), biofilm formation (4 of 37), pathogen suppression (15 of 37) or biomass production (2 of 37) (*SI Appendix*, Fig. S2). Moreover, consortia performance did not show clear relationship with increasing consortia richness regarding any of the measured traits (*SI Appendix*, Fig. S3). We also tested if the consortia performance could be predicted based on the sum of monoculture performance of individual mutants, assuming that mutant performance is not affected by interactions between the consortia members. Only one significant positive relationship was found between the predicted consortia suppressiveness, and the size of the inhibition halo observed *in vitro* lab measurements (Fig. 2A-D). This suggest that consortia functioning measured *in vitro* was poorly predicted by the sum of individual member performances except for the pathogen suppression.

**Figure 2.**
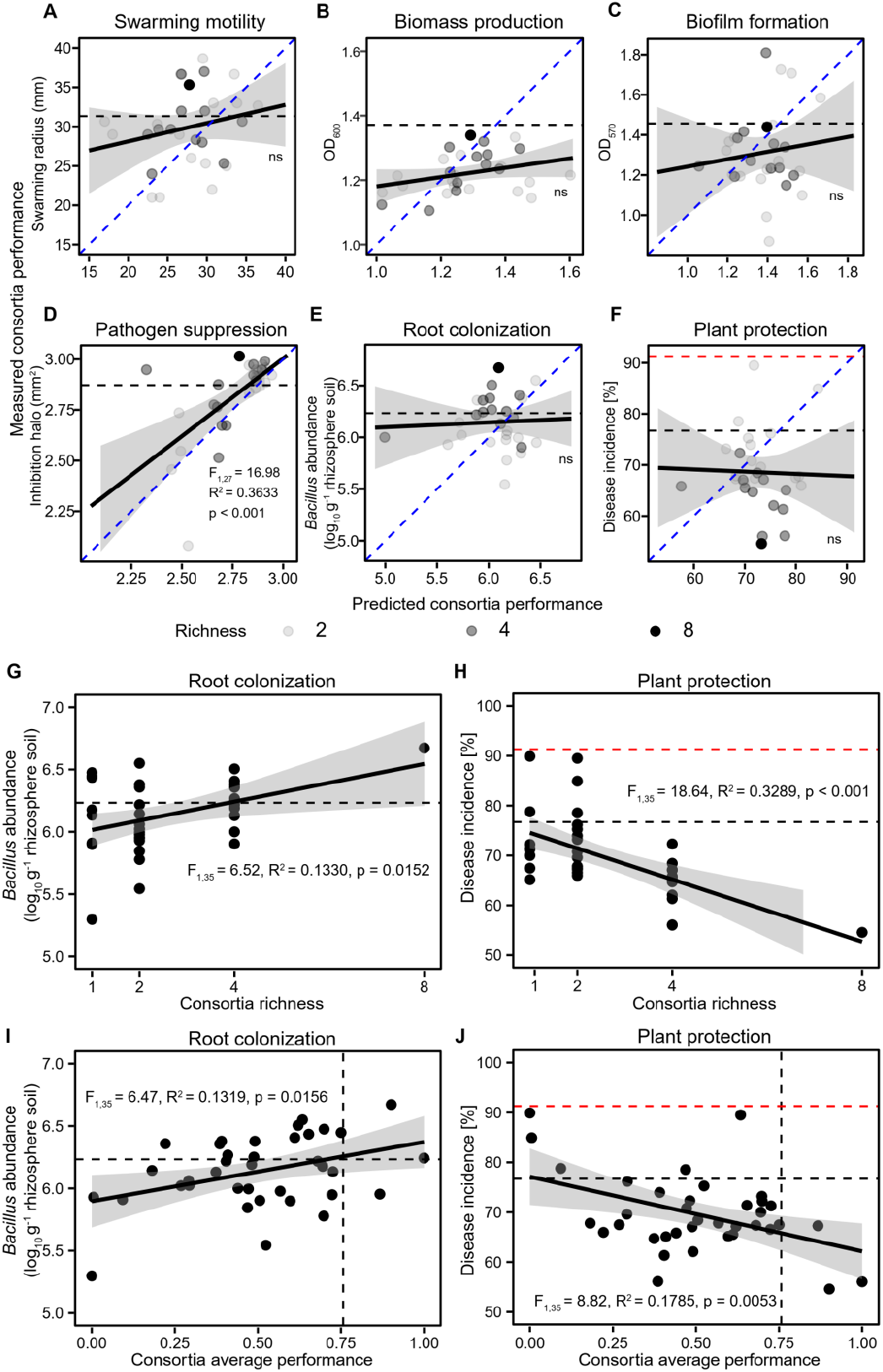
The relationships between predicted and observed consortia performance measured *in vitro* and correlations between plant performance, consortia richness and consortia average performance measured *in vivo*. Panels A-F, show correlations between predicted and observed consortia performance regarding swarming motility, biomass production, biofilm formation, pathogen suppression, root colonization and plant protection, respectively (blue dashed lines show 1:1 theoretical fit and solid black lines show the fitted regression between predicted and observed values). Panels G and H show regression models where root colonization and plant protection were explained by *B. amyloliquefaciens* T-5 consortia richness, respectively. Panels I and J show regression models where root colonization and plant protection were explained by *B. amyloliquefaciens* T-5 consortia average performance measured *in vitro*, respectively. In all panels, the black dashed lines show the performance of the wild-type strain, while red dashed lines in panels F, H and J show the disease incidence of pathogen-only control treatment (ns denotes for non-significant relationship; ** denotes for statistical significance at p < 0.01; * denotes for statistical significance at p < 0.05).

We next tested how consortia root colonization and plant protection *in vivo* was affected by the consortia diversity. While only 7 of 37 of consortia showed improved rhizosphere colonization, around half of them (18 of 37) exhibited a clear increase in plant protection compared to the wild-type strain (at 30 dpi, *SI Appendix*, Fig. S4). While root colonization or plant protection could not be predicted based on the sum of performance of individual mutants measured *in vitro* (Fig. 2E-F), both root colonization (Fig. 2G, F _1,35_ = 6.52, R^2^ = 0.1330, p = 0.0152) and plant protection (Fig. 2H, F _1,35_ = 18.64, R^2^ = 0.3289, p < 0.001) improved with increasing consortia richness, and were positively correlated with the consortia average performance measured *in vitro*, indicative of positive diversity-ecosystem functioning relationship (root colonization: F _1,35_ = 6.47, R^2^ = 0.1319, p = 0.0156; plant protection: F _1,35_ = 8.82, R^2^ = 0.1786, p = 0.0053; Fig. 2I-J). Finally, we analyzed the significance of mutant identity effects on the consortia performance *in vivo*. The presence of M54 mutant (efficient in biofilm formation) significantly increased consortia root colonization, while the presence of mutants M59 (efficient in biomass production) and M143 (efficient in biofilm formation) was significantly associated with improved plant protection (SI Appendix, Fig. S5, Table S8). Crucially, the effect of consortia richness remained significant after sequential removal of each mutant and refitting of the model, which demonstrates that the effect of diversity was relatively more important compared to mutant identity effects (*SI Appendix*, Table S9). Together, these data suggest that mutant consortia diversity was positively linked with consortia performance *in vivo*, which was associated with consortia mean performance and pathogen suppression measured *in vitro*.

### The relative effect of different mutant traits on rhizosphere colonization and plant protection varies temporally during the plant growth

To explore potential underlying mechanisms between consortia diversity and improved performance, we focused on analyzing the dynamics of root colonization and plant protection using 47 individual *B. amyloliquefaciens* T-5 mutants at 5, 15 and 30 dpi time points. Specifically, we tested the insurance hypothesis (87), which predicts that a community composed of functionally diverse genotypes is likely to perform better because of the likelihood that some mutants will thrive as prevailing conditions change during the plant growth, providing increased stability for the plant-microbe interaction. Trait correlation with the root colonization and plant protection became more significant with time and most significant correlations were observed at the final time point (30 dpi, followed by middle (15 dpi) and early (5 dpi) time points (Table 1, *SI Appendix*, Figure S5). Specifically, high swarming motility predicted the rhizosphere colonization during the seedling stage (5 dpi, Table 1; Fig 3A, F_1,46_ = 7.65, R^2^ = 0.1239, p = 0.0082), while swarming motility was positively associated with improved plant protection at the flowering stage (30 dpi, Table 1; Fig 3F, F_1,46_ = 15.08, R^2^ = 0.2306, p < 0.001). Similarly, biofilm formation had positive associations with root colonization (Table 1; Fig 3C, 15dpi: F_1,46_ = 4.40, R^2^ = 0.0675, p = 0.0416, 30dpi: F_1,46_ = 7.62, R^2^ = 0.1209, p = 0.0089) and plant protection during vegetative and flowering stages at 15 and 30 dpi, respectively (Table 1; Fig 3H, 15dpi: F_1,46_ = 6.44, R^2^ = 0.1038, p = 0.0146, 30dpi: F_1,46_ = 8.69, R^2^ = 0.1406, p = 0.0050). Pathogen suppression was positively associated with root colonization and plant protection at 30 dpi (Table 1; Fig 3D, F_1,46_ = 7.65, R^2^ = 0.1239, p = 0.0082; Fig 3I, F_1,46_ = 15.65, R^2^ = 0.2538, p < 0.001), while biomass production was not significantly associated with either root colonization or plant protection at any time points (Table 1; Fig 3B, 3G). As a result, the mean performance of mutants (‘Monoculture average performance’ index based on mean of all traits) was significantly correlated with both root colonization (Fig. 3E, F_1,46_ = 7.77, R^2^ = 0.1259, p = 0.0077) and plant protection (Fig. 3J, F_1,46_ = 28.26, R^2^ = 0.3671, p < 0.001) at the flowering stage (30 dpi). Together, these results suggest that while mutants with high trait values in biofilm formation, swarming motility and pathogen suppression had positive effects on root colonization and plant protection, their relative importance varied depending on the growth stage of the plant. Inclusion of multiple specialized mutants hence likely improved the consortia functioning via insurance effects by providing more stable and robust performance throughout the plant growth cycle.

**Table 1.** ANOVA table summarizing the effects of mutant traits measured *in vitro* on the root colonization and plant protection. Separate models were run for each dependent variable at different time points (5, 15, and 30 dpi) and all response variables were treated as continuous variables (bacterial abundances were log-transformed before the analysis). Table data represent only the most parsimonious models based on the Akaike’s information criterion (AIC) where ‘NA’ denotes variables that were not retained in the final models, ‘df’ denotes degrees of freedom and ‘R^2^’ denotes total variance explained by regression coefficient of determination. The arrows represent the direction of coefficient values: ↑: coefficient > 0; ↓: coefficient < 0. Significant effects (p < 0.05) are highlighted in bold.

| Mutant trait | Day post pathogen inoculation (dpi) |  |  |  |  |  |  |  |  |
| --- | --- | --- | --- | --- | --- | --- | --- | --- | --- |
|  | 5 dpi |  |  | 15 dpi |  |  | 30 dpi |  |  |
|  | Df | F | P | Df | F | P | Df | F | P |
| Root colonization (*Bacillus* abundance – log CFU g<sup>-1</sup> rhizosphere soil) |  |  |  |  |  |  |  |  |  |
| Swarming motility | 1 | 5.32 | 0.0259↑ | 1 | 0.006 | 0.9404↑ | 1 | 0.008 | 0.9302↑ |
| Biomass production | 1 | 0.46 | 0.5032↓ | 1 | 0.58 | 0.4512↑ | 1 | 0.07 | 0.7886↓ |
| Biofilm formation | 1 | 0.69 | 0.4103↑ | 1 | 4.34 | 0.0433↑ | 1 | 5.64 | 0.0220↑ |
| Pathogen suppression | 1 | 2.04 | 0.1609↓ | 1 | 0.006 | 0.9410↓ | 1 | 8.10 | 0.0068↑ |
| No. of residuals | 43 | R<sup>2</sup> = 0.0875 |  | 43 | R<sup>2</sup> = 0.0193 |  | 43 | R<sup>2</sup> = 0.1730 |  |
|  |  | AIC: 16.18 |  |  | AIC: 3.82 |  |  | AIC: 25.06 |  |
| Plant protection (Disease incidence [%]) |  |  |  |  |  |  |  |  |  |
| Swarming motility |  | NA | NA | 1 | 0.34 | 0.5620↑ | 1 | 6.89 | 0.0119↓ |
| Biomass production |  | NA | NA | 1 | 0.94 | 0.3369↓ | 1 | 0.23 | 0.6339↑ |
| Biofilm formation |  | NA | NA | 1 | 5.71 | 0.0213↓ | 1 | 12.34 | 0.0016↓ |
| Pathogen suppression |  | NA | NA | 1 | 0.14 | 0.7090↓ | 1 | 20.90 | <0.001↓ |
| No. of residuals |  | NA |  | 43 | R<sup>2</sup> = 0.0625 |  | 43 | R<sup>2</sup> = 0.4294 |  |
|  |  | NA |  |  | AIC: -101.24 |  |  | AIC: -112.38 |  |

**Figure 3.**
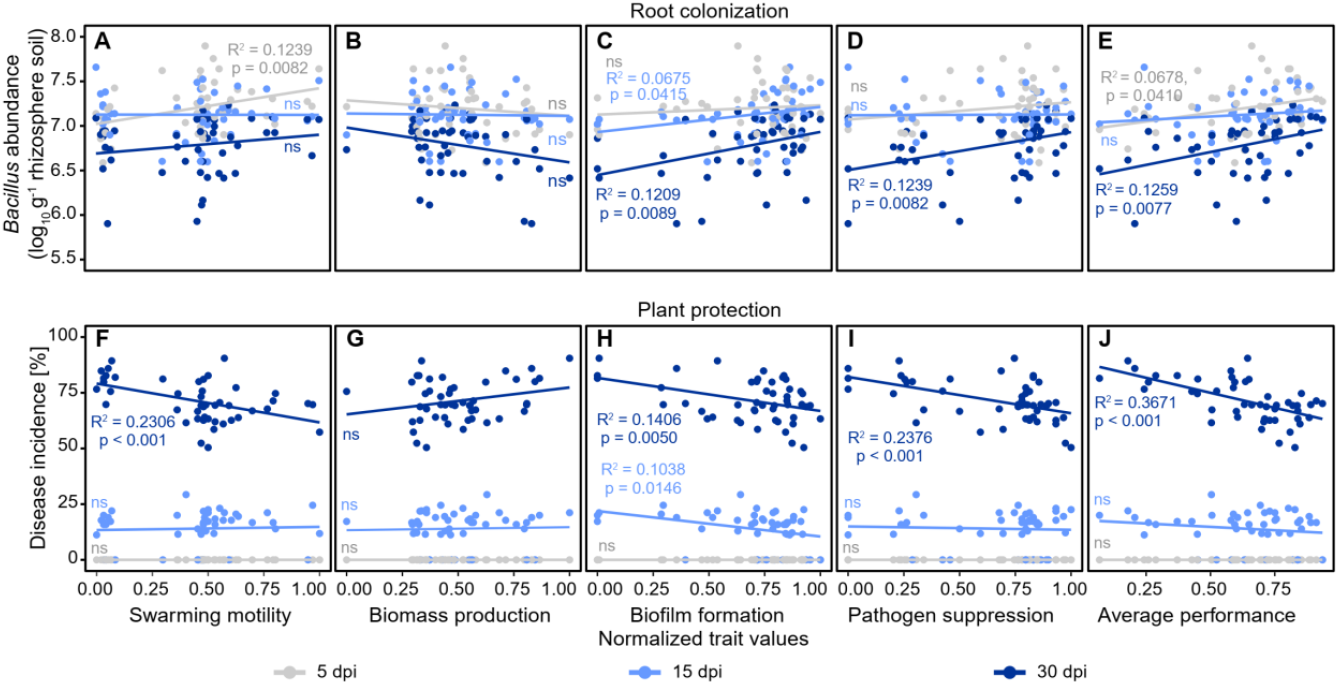
Regression analysis explaining root colonization and plant protection of 47 *B. amyloliquefaciens* T-5 mutants based on their trait values measured *in vitro* at different sampling time points. Panels A-E and F-J show root colonization and plant protection, respectively, correlated with different traits at 5 dpi (grey), 15 dpi (light blue) and 30 dpi (dark blue) time points. Significant relationships and R-squared values are shown in panels with colors corresponding to the sampling time points (‘ns’ denotes for non-significant relationship).

## Discussion

In this work, we tested if increasing intra-species diversity of *B. amyloliquefaciens* T-5 bacterium via mutagenesis could offer a viable strategy for improving mutant consortia multifunctionality and plant growth-promotion. Our results show that mutations that improved bacterial performance regarding one trait often led to specialism and reduced performance regarding other traits. Such trait trade-offs experienced at the individual genotype level could be overcome by assembling consortia of specialist mutants, that showed increase in average trait performance. Crucially, the consortia richness and average trait performance correlated positively with increased root colonization and plant protection, indicative of increased consortia multifunctionality and improved plant health. Together, these findings suggest that increasing intra-species functional diversity could offer an easy solution for improving the performance of bacterial communities without introducing potential inter-species conflicts.

We specifically focused on four *B. amyloliquefaciens* traits that have previously been linked to plant growth-promotion: swarming motility, biomass production, biofilm formation and direct pathogen suppression via antibiosis (69–72). While most insertions reduced the strains’ performance overall, several of them were associated with improvements regarding at least one measured trait. However, all trait improvements were costly and reduced mutant performances regarding other traits, indicative of trade-offs (88). Such costs of adaptation are common with microbes (89) and have previously been linked to a wide range of functions, including metabolism, antagonism, motility and stress resistance (66, 90). Overall, transposon insertions were identified in several genes associated with broad range of functions (*SI Appendix*, Figure S1, Dataset S2). With eight specialist strains that were used for consortia assembly experiment, increased swarming motility was associated with insertions in *parE* (DNA topoisomerase IV subunit B) and *DeoR* (DNA-binding transcriptional repressor) genes instead of insertions in flagellar genes. The *parE* gene has previously been linked to antibiotic resistance (91), while *DeoR* is known to repress dra-nupC-pdp operon, which encodes three enzymes required for deoxyribonucleoside and deoxyribose utilization (92). While it remains unclear how these genes were linked with swarming motility, their disruption affected also other traits as evidenced by reduced biofilm formation. Increased biomass production was linked to disruption of *comQ* (competence protein) and *hutI* (imidazolonepropionase) genes and trade-offs with the other three measured traits. *ComQ* gene controls the production of ComX pheromone, which regulates quorum sensing in density-dependent manner in *Bacillus* (93) Mutation in this gene could have thus led to a loss of response to crowding, resulting in higher biomass production. Moreover, *ComQ* gene has recently been linked to antimicrobial activity (94), which could explain reduction in the pathogen suppression by this mutant. Insertion in histidine utilization (hut) system gene, *hutI*, could have potentially impaired catabolite and amino acid repression, resulting in improved biomass production with one of the mutants (95, 96). Increased biofilm formation was also linked to insertion in hut system (*hutU*) with one mutant, while the other mutant had insertion in *YsnB* gene, which encodes for a putative metallophosphoesterase. While both insertions were linked to trade-offs with swarming motility and biomass production, they are not commonly associated with biofilm formation in *Bacillus* (18, 70). Finally, moderate improvement in pathogen suppression was observed with two mutants that had insertions in *nhaC* (sodium-proton antiporter) and *dfnG* (difficidin polyketide synthesis) genes. The *nhaC* gene is known to act as repressor for Pho regulon in *Bacillus*, which controls response to phosphate limitation via production of alkaline phosphatases (97). Interestingly, mutations in Pho regulon have also been linked to increased antibiotics production with several *Streptomyces* species via unknown mechanisms (98). The *dfnG* gene controls the production of difficidin antibiotic, which has previously been linked to biocontrol activity against fire blight (99) and *Xanthomonas oryzae* rice pathogen (100) and could have hence also had a positive effect on *R. solanacearum* suppression. Similar to the other specialist mutants, insertions associated with increased pathogen suppression led to trade-offs with other measured traits. While more work is required to link these mutations with associated traits at the molecular level in the future, our findings show that all above trait improvements achieved via mutagenesis resulted in trade-offs with other traits.

To overcome trait trade-offs experienced at the individual mutant level, we tested if we could improve the plant growth-promotion by combining individual specialist mutants into multifunctional consortia. We found that mutant performance measured in monocultures was a poor predictor of *B. amyloliquefaciens* performance in coculture consortia, except for the pathogen suppression. This suggest that while the selected mutants did not show direct antagonism towards each other, they likely interacted in other ways leading to unpredictable trait expression when embedded in consortia (for example via certain emergent effects (101)). Despite this, we found clear diversity effects where consortia richness and average performance were positively associated with both plant root colonization and protection from *R. solanacearum* pathogen infection. This suggests that consortia that performed well regarding all measured traits on average had improved capacity to colonize rhizosphere and suppress the pathogen. While similar positive diversity-ecosystem functioning relationships have previously been found in more complex *Pseudomonas* (6, 34, 37), leaf bacterial (102) and grass-land soil microbiomes (103), we here show that this pattern also holds along with intra-species diversity gradient. While important mutant identity effects were also observed, omission of these strains did not change the significance of underlying diversity effects, highlighting the importance of interactions between the consortia members in determining the positive effects on the plant health.

Positive diversity effects have previously been explained by facilitation, ecological complementarity, and division of labor, which can reduce competition between the consortia members within or between niches (18, 19, 28). Moreover, high diversity could provide stability for consortia functioning via insurance effects by increasing the likelihood that certain members survive in the soil after the inoculation (63, 87). While our experiments were not designed to disentangle the relative importance of these potential mechanisms, we analyzed which mutant traits could significantly explain the dynamics of root colonization and plant protection at seedling, vegetative and flowering stages of the tomato growth by focusing on 47 mutants. While the effect of *Bacillus* biomass production was consistently non-significant, both motility and biofilm formation were positively associated with the root colonization. However, motility was significant only at the seedling stage, while biofilm became significant during vegetative and flowering stages. In line with ecological succession often taking place in the rhizosphere (69), high motility might have allowed faster colonization of relatively ‘sterile’ young roots by *Bacillus*, while biofilm formation could have promoted stress tolerance and resource competition in more diverse and mature microbial communities during the later stages of tomato growth (104, 105). Interestingly, increase in pathogen suppression, biofilm formation and motility were positively associated with improved plant protection at the flowering stage, which suggests that all these traits were positively associated with the ecosystem functioning in terms of plant health. It is thus possible that consortia were together able to overcome the trait trade-offs experienced at the individual mutant level, leading to improved and more stable ecosystem functioning in time. Future studies addressing genotype frequencies and population dynamics in space and time are however required to test if such stability rises due to ecological complementarity, insurance effects or population asynchrony (106).

In conclusion, we here demonstrate that the beneficial effects provided by a single *B. amyloliquefaciens* bacterium can be improved via combination of synthetic biology (transposon mutagenesis) and community assembly, leading to improved consortia multifunctionality. Our approach highlights the importance of intra-species genetic diversity for the ecosystem functioning and provides a trait-based approach for designing microbial communities for biotechnological applications. Moreover, our approach does not require a priori knowledge on specific genes or molecular mechanisms, but instead relies on generation of trait variation which is screened and selected for by the experimenter and can also help in identify novel functional roles of previously characterized genes. While the benefit of this method was here demonstrated in the context of agriculture, it could be applied in other biotechnological contexts, including biofermentation, waste degradation and food manufacturing. Future work focusing on the population dynamics, metabolism and gene expression of different mutants in complex consortia will help to understand the relative importance of ecological complementarity, division of labor and facilitation for the consortia ecosystem functioning.

## Acknowledgments

We thank Daniel Rozen, Benoit Stenuit and Zheren Zhang for valuable comments on the manuscript. This research was supported by the National Key Research and Development Program of China (2021YFD1900100), the National Natural Science Foundation of China (42090064, 41922053 and 31972504), the Fundamental Research Funds for the Central Universities (KY2201719, KYT201802). V-PF is supported by the Royal Society Research Grants (RSG\R1\180213 and CHL\R1\180031) and jointly by a grant from UKRI, Defra, and the Scottish Government, under the Strategic Priorities Fund Plant Bacterial Diseases programme (BB/T010606/1) at the University of York.

---

**Table S1.** Bacterial strains and plasmid used in this study.

| Strain or plasmid | Characteristics | Source |
| --- | --- | --- |
| *Ralstonia solanacearum*<br>QL-Rs1115 | Causative agent of Bacterial wilt | Z. Wei *et al.* 2011 (64) |
| *Bacillus amyloliquefaciens* T-5 | Rhizosphere isolate capable of suppressing the growth of *Ralstonia solanacearum* | S. Y. Tan *et al.* 2013 (50),<br>X. F. Wang *et al.* 2017<br>(107) |
| pMarA plasmid | promoter σA, kmR ApR EmR pUC19 carrying TnYLB-1 transposon, mariner-Himar1 transposase | Y. Le Breton *et al.* 2006<br>(52) |

**Table S2.**
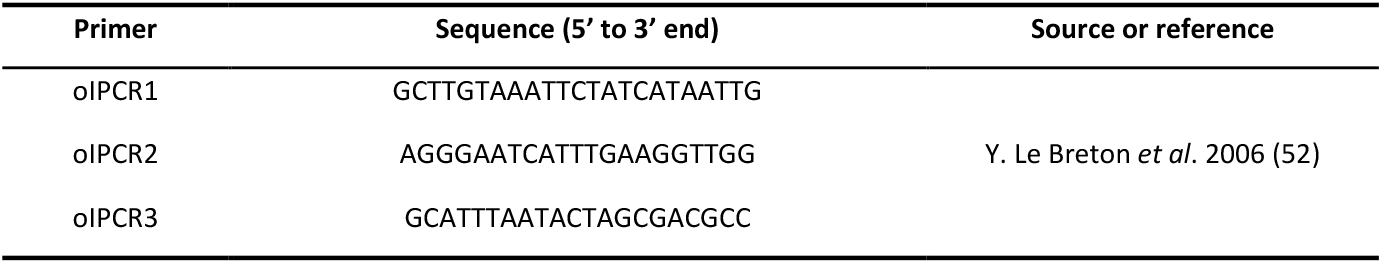
Primers used in this study.

**Table S3.** Phenotypic characteristics of the eight *B. amyloliquefaciens* mutants used for assembly of consortia. Blue, red and grey cells show increase, decrease or no change (ns) in normalized trait values relative to the wild-type strain (ANOVA followed by Two-sided Dunnett’s Multiple Comparisons at p < 0.05; see details in Table S4). The rightmost column (*) shows the functional categories (based on clusters of orthologous genes; COG) of different genes in parentheses, where capital letters denote for: C: Energy production and conversion; E: Amino acid transport and metabolism; R: General function prediction only; Q: Secondary metabolites biosynthesis, transport and catabolism; L: Replication, recombination and repair; K: Transcription.

| Mutant strain | Phenotypic characteristics |  |  |  | Disrupted gene | Annotated gene function (General function*) |
| --- | --- | --- | --- | --- | --- | --- |
|  | Swarming | Biomass | Biofilm | Pathogen suppression |  |  |
| M38 | ns | 0.92 | 0.90 | 1.02 | *nhaC* | sodium-proton antiporter (C) |
| M54 | ns | 0.89 | 1.11 | ns | *hutU* | histidine utilization (E) |
| M59 | 0.61 | 1.13 | 0.85 | 0.73 | *comQ* | competence protein (R) |
| M78 | 0.48 | 0.63 | ns | 1.03 | *dfnG* | difficidin polyketide synthase (Q) |
| M108 | 1.21 | ns | 0.89 | ns | *parE* | DNA topoisomerase IV subunit B (L) |
| M109 | 0.86 | 1.21 | 0.81 | ns | *hutI* | imidazolonepropionase (Q) |
| M124 | 1.30 | ns | 0.83 | 0.97 | *DeoR* | DNA-binding transcriptional repressor (K) |
| M143 | 0.67 | 0.85 | 1.26 | ns | *YsnB* | metallophosphoesterase (R) |

**Table S4.** Comparison of trait values of eight mutants selected to assemble consortia relative to the wild-type strain based on Two-sided Dunnett’s Multiple Comparisons. Significant differences are shown on bold and arrows show the increase (upwards) and decrease (downwards) in trait values.

| Strain | P value |  |  |  |
| --- | --- | --- | --- | --- |
|  | Swarming | Biomass | Biofilm | Pathogen suppression |
| M38 | 0.8683 | **0.0057 ↓** | **0.0023 ↓** | **0.0173 ↑** |
| M54 | 0.1451 | **< 0.001 ↓** | **< 0.001 ↑** | 0.9653 |
| M59 | **< 0.001 ↓** | **< 0.001 ↑** | **< 0.001 ↓** | **< 0.001 ↓** |
| M78 | **< 0.001 ↓** | **< 0.001 ↓** | 0.6918 | **< 0.001 ↑** |
| M108 | **< 0.001 ↑** | 0.0747 | **< 0.001 ↓** | 0.9998 |
| M109 | **0.0022 ↓** | **< 0.001 ↑** | **< 0.001 ↓** | 1.0000 |
| M124 | **< 0.001 ↑** | 0.1594 | **< 0.001 ↓** | **0.0020 ↓** |
| M143 | **< 0.001 ↓** | **< 0.001 ↓** | **< 0.001 ↑** | 0.9998 |

**Table S5.** Composition of the *B. amyloliquefaciens* consortia consisting of 8 mutants of *B. amyloliquefaciens* T-5 (named as M38, M54, ME59, M78, M108, M109, M124, and M143, respectively). 1 and 0 in the table denotes that the presence and absence of the corresponding mutant in the community, respectively.

| Consortia<br>code | Consortia members |  |  |  |  |  |  |  | Consortia<br>richness |
| --- | --- | --- | --- | --- | --- | --- | --- | --- | --- |
|  | M38 | M54 | M59 | M78 | M108 | M109 | M124 | M143 |  |
|  | Pathogen<br>suppression+ | Biofilm<br>formation+ | Biomass<br>production+ | Pathogen<br>suppression+ | Swarming<br>motility+ | Biomass<br>production+ | Swarming<br>motility+ | Biofilm<br>formation+ |  |
| 1 | 1 | 0 | 0 | 0 | 0 | 0 | 0 | 0 | 1 |
| 2 | 0 | 1 | 0 | 0 | 0 | 0 | 0 | 0 | 1 |
| 3 | 0 | 0 | 1 | 0 | 0 | 0 | 0 | 0 | 1 |
| 4 | 0 | 0 | 0 | 1 | 0 | 0 | 0 | 0 | 1 |
| 5 | 0 | 0 | 0 | 0 | 1 | 0 | 0 | 0 | 1 |
| 6 | 0 | 0 | 0 | 0 | 0 | 1 | 0 | 0 | 1 |
| 7 | 0 | 0 | 0 | 0 | 0 | 0 | 1 | 0 | 1 |
| 8 | 0 | 0 | 0 | 0 | 0 | 0 | 0 | 1 | 1 |
| 9 | 1 | 1 | 0 | 0 | 0 | 0 | 0 | 0 | 2 |
| 10 | 0 | 1 | 1 | 0 | 0 | 0 | 0 | 0 | 2 |
| 11 | 1 | 0 | 0 | 1 | 0 | 0 | 0 | 0 | 2 |
| 12 | 0 | 1 | 0 | 0 | 1 | 0 | 0 | 0 | 2 |
| 13 | 1 | 0 | 0 | 0 | 0 | 0 | 1 | 0 | 2 |
| 14 | 0 | 0 | 1 | 1 | 0 | 0 | 0 | 0 | 2 |
| 15 | 1 | 0 | 0 | 0 | 0 | 0 | 0 | 1 | 2 |
| 16 | 0 | 0 | 1 | 0 | 0 | 1 | 0 | 0 | 2 |
| 17 | 0 | 0 | 0 | 1 | 1 | 0 | 0 | 0 | 2 |
| 18 | 0 | 0 | 1 | 0 | 0 | 0 | 1 | 0 | 2 |
| 19 | 0 | 1 | 0 | 0 | 0 | 1 | 0 | 0 | 2 |
| 20 | 0 | 0 | 0 | 1 | 0 | 0 | 0 | 1 | 2 |
| 21 | 0 | 0 | 0 | 0 | 1 | 0 | 0 | 1 | 2 |
| 22 | 0 | 0 | 0 | 0 | 1 | 1 | 0 | 0 | 2 |
| 23 | 0 | 0 | 0 | 0 | 0 | 1 | 1 | 0 | 2 |
| 24 | 0 | 0 | 0 | 0 | 0 | 0 | 1 | 1 | 2 |
| 25 | 1 | 1 | 0 | 1 | 1 | 0 | 0 | 0 | 4 |
| 26 | 0 | 1 | 0 | 1 | 0 | 1 | 0 | 1 | 4 |
| 27 | 1 | 0 | 0 | 1 | 1 | 1 | 0 | 0 | 4 |
| 28 | 0 | 1 | 0 | 0 | 1 | 0 | 1 | 1 | 4 |
| 29 | 1 | 1 | 1 | 1 | 0 | 0 | 0 | 0 | 4 |
| 30 | 0 | 1 | 0 | 0 | 0 | 1 | 1 | 1 | 4 |
| 31 | 1 | 0 | 0 | 0 | 1 | 1 | 1 | 0 | 4 |
| 32 | 0 | 1 | 1 | 0 | 1 | 0 | 0 | 1 | 4 |
| 33 | 1 | 0 | 1 | 0 | 0 | 1 | 1 | 0 | 4 |
| 34 | 0 | 0 | 1 | 1 | 0 | 1 | 0 | 1 | 4 |
| 35 | 1 | 0 | 1 | 1 | 0 | 0 | 1 | 0 | 4 |
| 36 | 0 | 0 | 1 | 0 | 1 | 0 | 1 | 1 | 4 |
| 37 | 1 | 1 | 1 | 1 | 1 | 1 | 1 | 1 | 8 |

**Table S6.** Comparison of *in vitro* traits between mutants belonging to three clusters using unpaired two-samples Wilcoxon test. Significant effects (p < 0.05) are highlighted on bold.

| Bacterial trait measured *in vitro* | Comparison | W | P |
| --- | --- | --- | --- |
| Swarming motility | Cluster1 (n=265) VS Cluster2 (n=112) | **11125** | **<0.001** |
|  | Cluster1 (n=265) VS Cluster3 (n=103) | **26874** | **<0.001** |
|  | Cluster2 (n=112) VS Cluster3 (n=103) | **11517** | **<0.001** |
| Biomass production | Cluster1 (n=265) VS Cluster2 (n=112) | **10869** | **<0.001** |
|  | Cluster1(n=265) VS Cluster3 (n=103) | **7754** | **<0.001** |
|  | Cluster2 (n=112) VS Cluster3 (n=103) | 4931 | 0.0664 |
| Biofilm formation | Cluster1 (n=265) VS Cluster2 (n=112) | **23061** | **<0.001** |
|  | Cluster1 (n=265) VS Cluster3 (n=103) | **18268** | **<0.001** |
|  | Cluster2 (n=112) VS Cluster3 (n=103) | **4238** | **<0.001** |
| Pathogen suppression | Cluster1 (n=265) VS Cluster2 (n=112) | **28464** | **<0.001** |
|  | Cluster1 (n=265) VS Cluster3 (n=103) | **27253** | **<0.001** |
|  | Cluster2 (n=112) VS Cluster3 (n=103) | **10300** | **<0.001** |

**Table S7.**
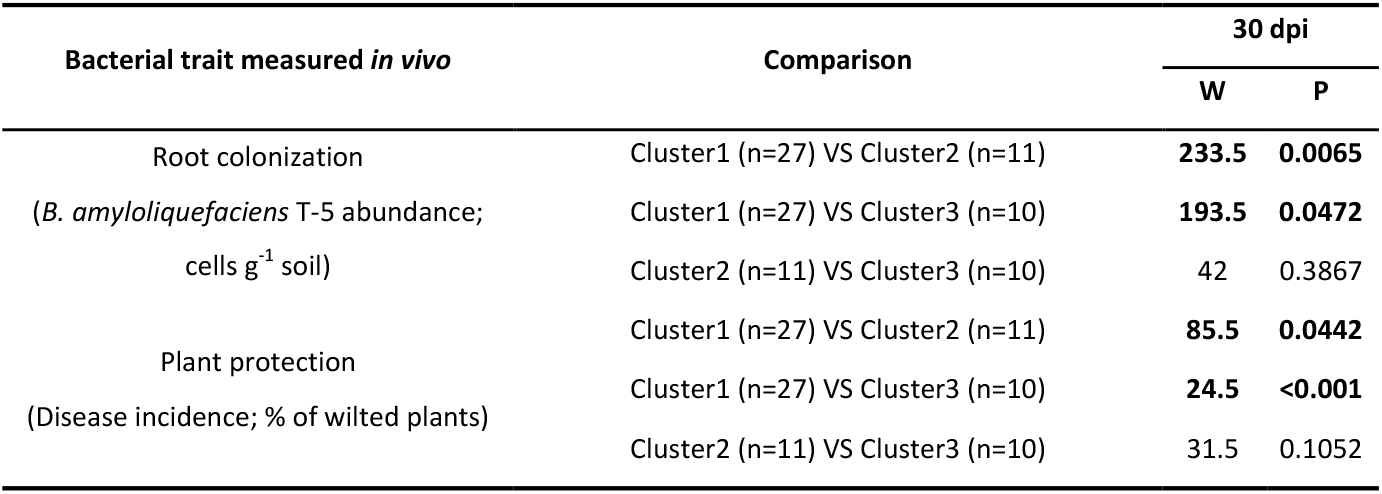
Comparison of root colonization and plant protection between mutants belonging to three clusters using unpaired two-samples Wilcoxon test. Significant effects (p < 0.05) are highlighted on bold and dpi denotes for days post-pathogen inoculation.

| Bacterial trait measured *in vivo* | Comparison | 30 dpi |  |
| --- | --- | --- | --- |
|  |  | W | P |
| Root colonization<br>(*B. amyloliquefaciens* T-5 abundance;<br>cells g<sup>-1</sup> soil) | Cluster1 (n=27) VS Cluster2 (n=11) | **233.5** | **0.0065** |
|  | Cluster1 (n=27) VS Cluster3 (n=10) | **193.5** | **0.0472** |
|  | Cluster2 (n=11) VS Cluster3 (n=10) | 42 | 0.3867 |
| Plant protection<br>(Disease incidence; % of wilted plants) | Cluster1 (n=27) VS Cluster2 (n=11) | **85.5** | **0.0442** |
|  | Cluster1 (n=27) VS Cluster3 (n=10) | **24.5** | **<0.001** |
|  | Cluster2 (n=11) VS Cluster3 (n=10) | 31.5 | 0.1052 |

**Table S8.** Comparison of mutant identity effects on consortia root colonization and plant protection in the absence and presence of each mutant based on unpaired two-samples Wilcoxon test. Significant effects (p < 0.05) are highlighted on bold.

| Mutant Strain ID | Phenotype | Comparison | Root colonization<br>(*Bacillus* abundance) |  | Plant protection<br>(Disease incidence) |  |
| --- | --- | --- | --- | --- | --- | --- |
|  |  |  | W | P | W | P |
| M38 | Pathogen suppression+ | Included (n=12) VS Excluded (n=25) | 197 | 0.1314 | 116 | 0.2815 |
| M54 | Biofilm formation+ | Included (n=12) VS Excluded (n=25) | **266** | **0.0143** | 128 | 0.4909 |
| M59 | Biomass production+ | Included (n=12) VS Excluded (n=25) | 192.5 | 0.1729 | **78** | **0.0188** |
| M78 | Pathogen suppression+ | Included (n=12) VS Excluded (n=25) | 139 | 0.7333 | 113 | 0.2401 |
| M108 | Swarming motility+ | Included (n=12) VS Excluded (n=25) | 154 | 0.9096 | 120 | 0.3435 |
| M109 | Biomass production+ | Included (n=12) VS Excluded (n=25) | 166.5 | 0.6037 | 105 | 0.1507 |
| M124 | Swarming motility+ | Included (n=12) VS Excluded (n=25) | 177 | 0.3899 | 97 | 0.0887 |
| M143 | Biofilm production+ | Included (n=12) VS Excluded (n=25) | 155 | 0.8839 | **78** | **0.0188** |

**Table S9.** Comparison of the mutant identity effects and consortia richness on root colonization and plant protection. Richness was fitted sequentially after mutant identity effects (presence or absence). Both response variables were treated as continuous variables and *Bacillus* abundance data was log-transformed before the analysis. Significant effects (p < 0.05) are highlighted on bold.

| Identity | Phenotype | Df | Root colonization (*Bacillus* abundance) |  | Plant protection (Disease incidence) |  |
| --- | --- | --- | --- | --- | --- | --- |
|  |  |  | F | P | F | P |
| M38 | Pathogen suppression+ | 1 | **4604.70** | **<0.001** | **652.86** | **<0.001** |
| M54 | Biofilm formation+ | 1 | **3413.15** | **<0.001** | **476.77** | **<0.001** |
| M59 | Biomass production+ | 1 | **2665.22** | **<0.001** | **433.88** | **<0.001** |
| M78 | Pathogen suppression+ | 1 | **1719.05** | **<0.001** | **269.91** | **<0.001** |
| M108 | Swarming motility+ | 1 | **2021.15** | **<0.001** | **272.12** | **<0.001** |
| M109 | Biomass formation+ | 1 | **1515.47** | **<0.001** | **215.33** | **<0.001** |
| M124 | Swarming motility+ | 1 | **1431.96** | **<0.001** | **185.12** | **<0.001** |
| M143 | Biofilm production+ | 1 | **1035.52** | **<0.001** | **126.80** | **<0.001** |
| Richness |  |  | **9.07** | **0.0048** | **22.24** | **<0.001** |
| Linear term |  | 1 |  |  |  |  |
| Error |  | 29 |  |  |  |  |

**Figure S1.**
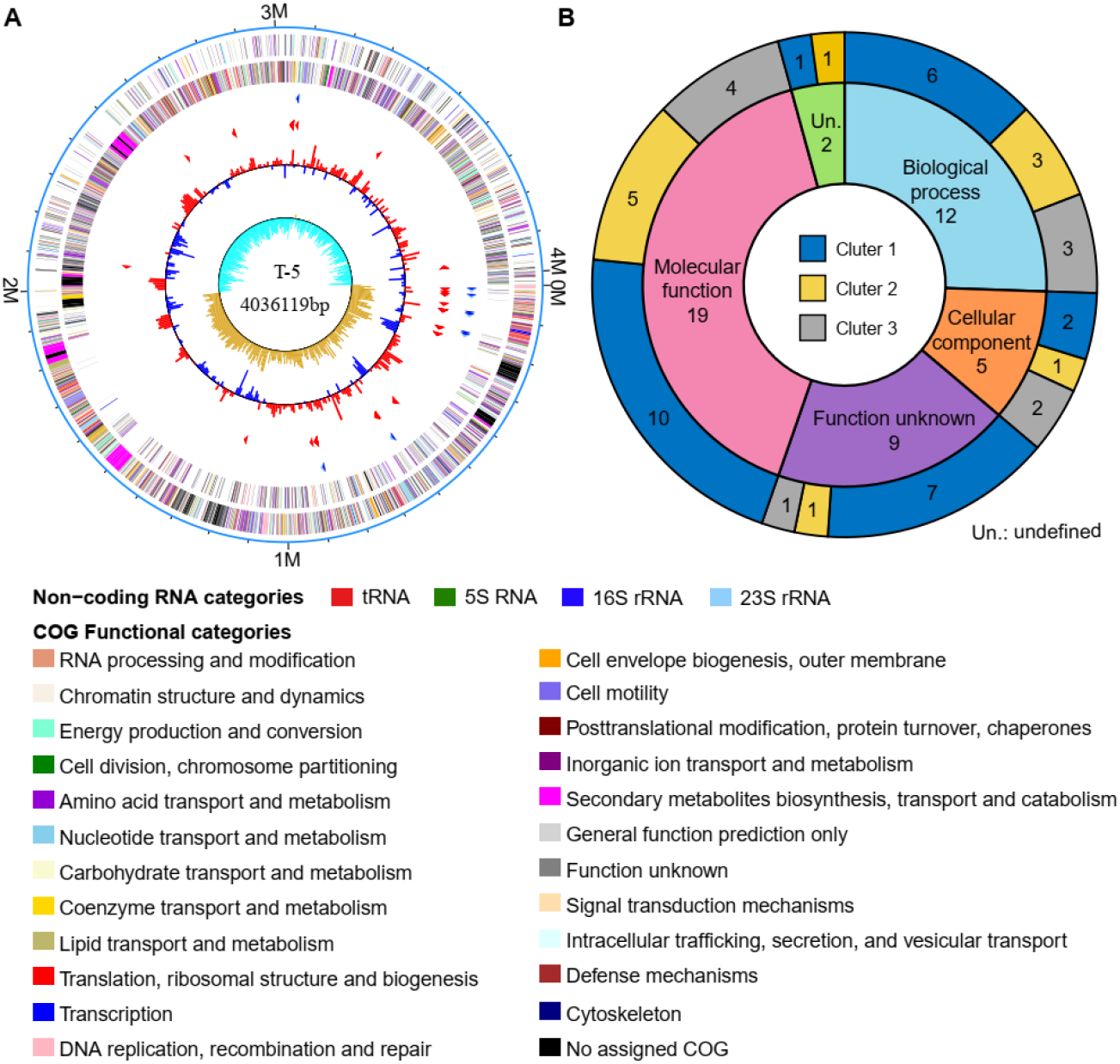
**A:** Graphical circular map of *Bacillus amyloliquefaciens* T-5 genome. Moving from the outside to the center different circles denote for: genes of the forward strand (colored by COG categories), genes of the reverse strand (colored by COG categories), RNA genes (tRNAs green, rRNAs red, other RNAs black), GC content and GC skew. **B:** The distribution of disrupted gene functions of mutants classified by each cluster based on the annotated reference genome of the *B. amyloliquefaciens* T-5 (detailed gene sequences of the disrupted genes is provided in Dataset S1).

**Figure S2.**
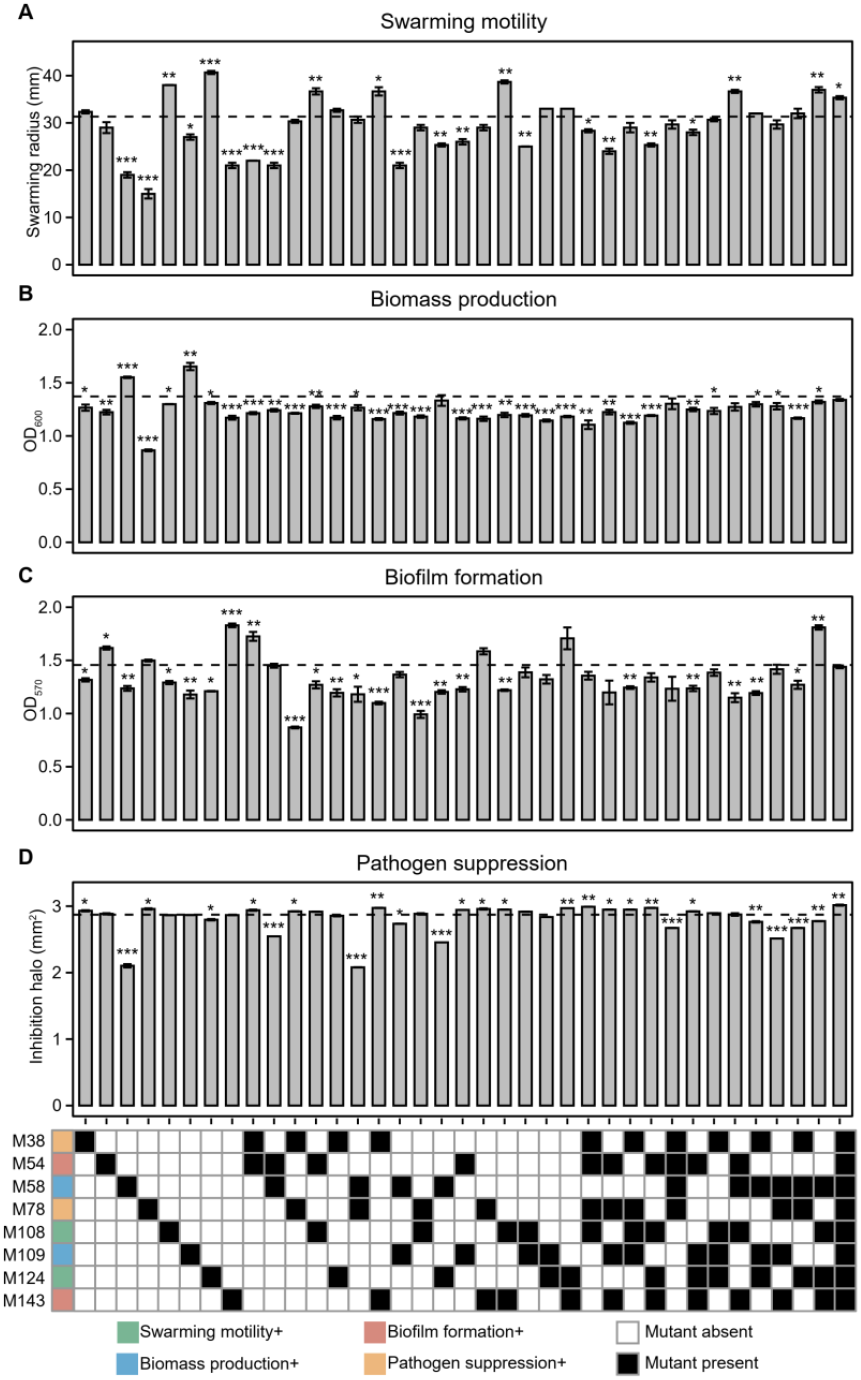
The performance of single mutants and mutant consortia relative to wild-type strain regarding four traits measured *in vitro*. (A) Swarming motility, (B) biomass production, (C) biofilm formation and (D) pathogen suppression. In panels A-D, the black dashed lines represent the performance of the wild-type strain and mean differences between consortia were analyzed using student’s t-test: *** denotes for statistical significance at p < 0.001; ** denotes for statistical significance at p < 0.01; * denotes for statistical significance at p < 0.05. Bottom panel shows the ‘trait specialism’ of eight mutants (colored squares) and their presence (black squares) and absence (white squares) in the tested consortia.

**Figure S3.**
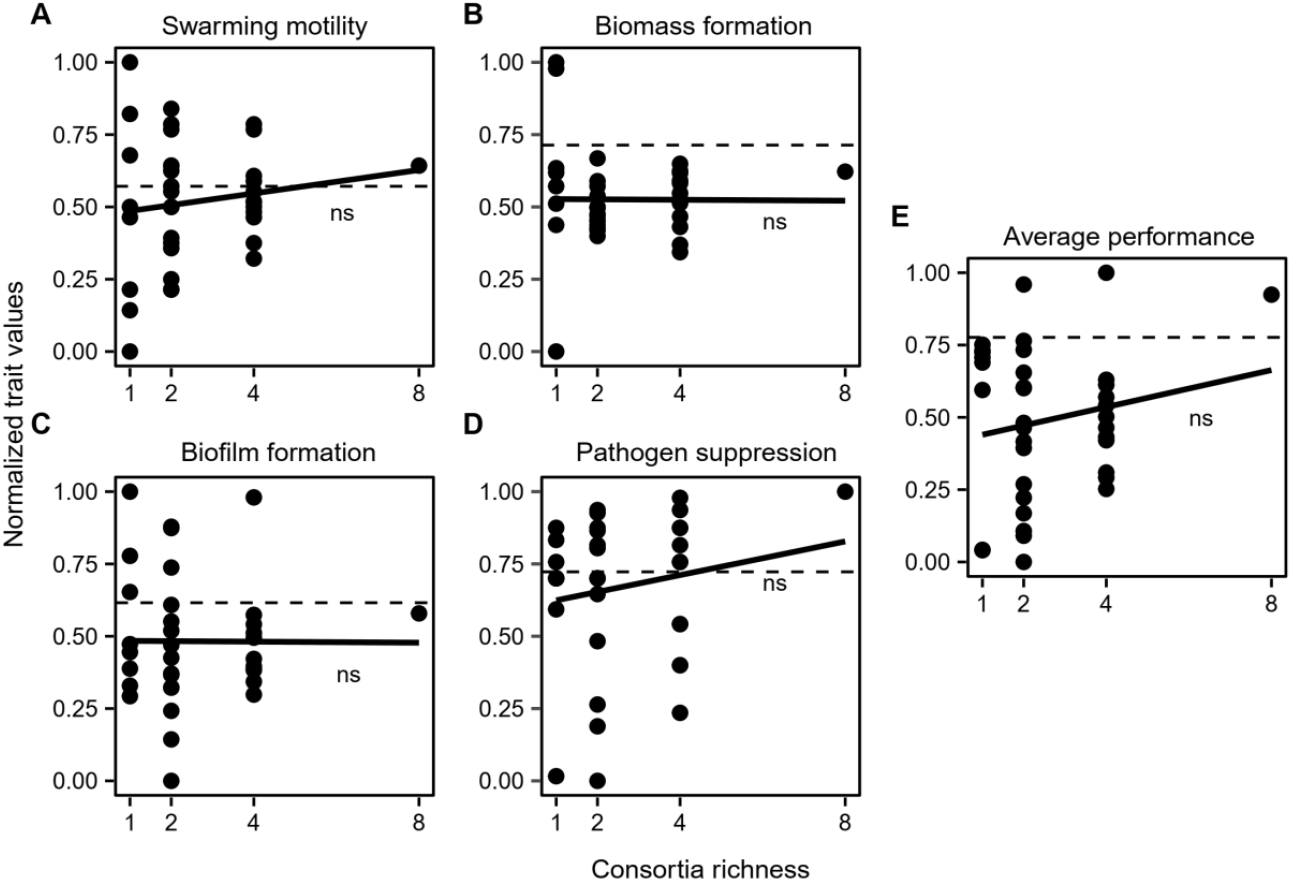
The relationship between *B. amyloliquefaciens* T-5 mutant consortia richness and measured consortia trait performance *in vitro*. The panels denote for swarming motility (A), biomass production (B), biofilm formation (C), and pathogen suppression (D) and consortia average performance (E; mean of all traits). In all panels, Y-axis show the normalized trait values, the black dashed line represents the performance of the wild-type strain and solid black line shows the fitted regression; ns denotes for non-significant relationship.

**Figure S4.**
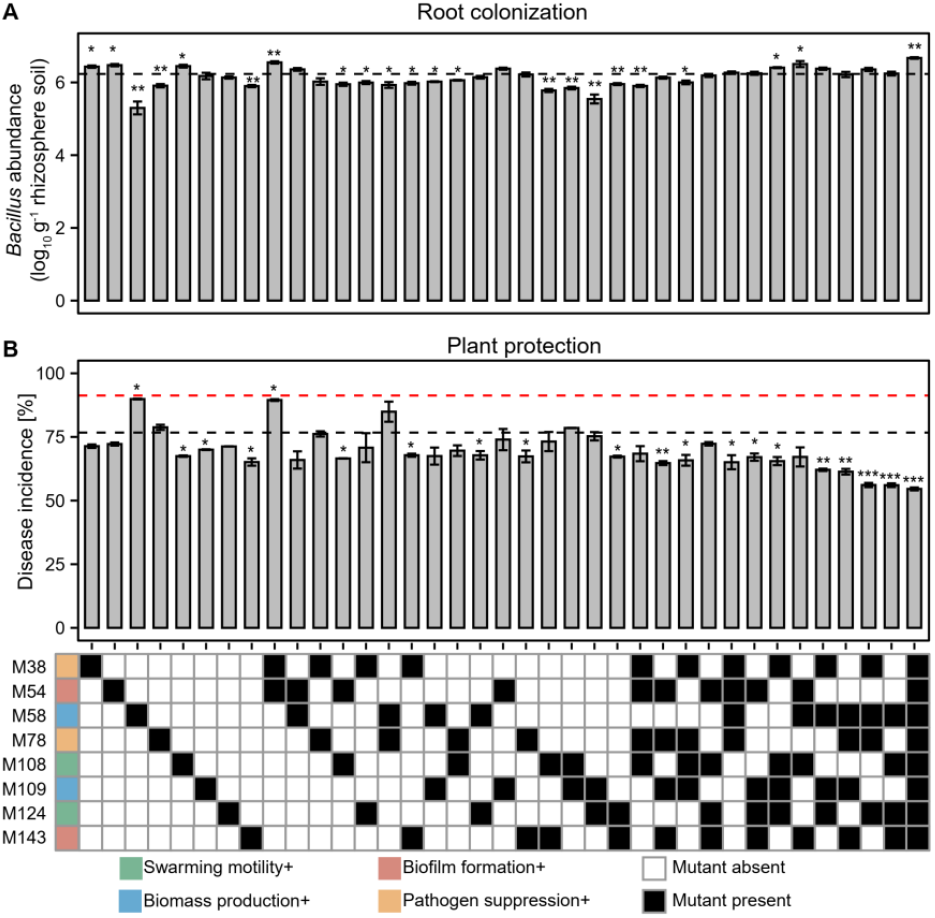
The performance of single mutants and mutant consortia relative to wild-type strain regarding root colonization and plant protection measured *in vivo*. Panel (A) shows root colonization and panel (B) plant protection. In both panels, the black and red dashed lines represent the performance of the wild-type strain and pathogen-only control treatments, respectively. Mean differences between consortia were analyzed using student’s t-test: *** denotes for statistical significance at p < 0.001; ** denotes for statistical significance at p < 0.01; * denotes for statistical significance at p < 0.05. Bottom panel shows the ‘trait specialism’ of eight mutants (colored squares) and their presence (black squares) and absence (white squares) in the tested consortia.

**Figure S5.**
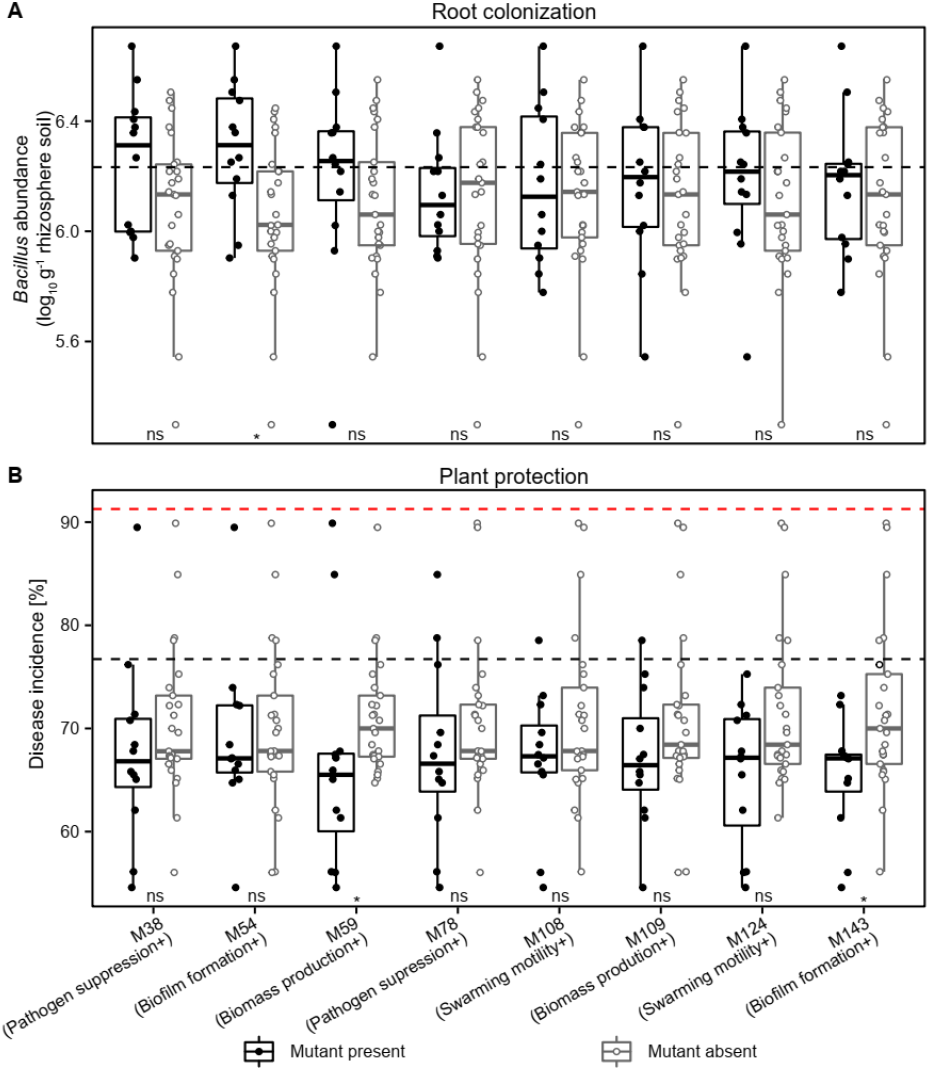
Analysis of mutant identity effects on consortia performance *in vivo*. The mutant identity effects were analyzed comparing consortia root colonization (A) and plant protection (B) in the absence and presence of each mutant. In both panels, the black dashed lines represent the performance of the wild-type strain, while red dashed line in panel (B) represents disease incidence in pathogen-only control treatment. The mutants’ ‘trait specialism’ is shown in parentheses on X-axis. Differences were analyzed using unpaired two-sample Wilcoxon test, where * denotes for statistical significance at p < 0.05; ns denotes for no significant (See details in SI Appendix, Table S7).

